# Spatial organization of nuclear pores in *Xenopus laevis* oocytes

**DOI:** 10.1101/2021.09.01.458492

**Authors:** Linda Ravazzano, Silvia Bonfanti, Roberto Guerra, Fabien Montel, Caterina A. M. La Porta, Stefano Zapperi

## Abstract

Nuclear pores are protein assemblies inserted in the nuclear envelope of eukaryotic cells, acting as main gates for communication between nucleus and cytoplasm. So far, nuclear pores have been extensively studied to determine their structure and composition, yet their spatial organization and geometric arrangement on the nuclear surface are still poorly understood. Here, we analyze super-resolution images of the surface of *Xenopus laevis* oocyte nuclei during development, and characterize the arrangement of nuclear pores using tools commonly employed to study the atomic structural and topological features of soft matter. To interpret the experimental results, we hypothesize an effective interaction among nuclear pores and implemented it in extensive numerical simulations of octagonal clusters mimicking typical pore shapes. Thanks to our simple model, we find simulated spatial distributions of nuclear pores that are in excellent agreement with experiments, suggesting that an effective interaction among nuclear pores exists and could explain their geometrical arrangement. Furthermore, our results show that the statistical features of the geometric arrangement of nuclear pores do not depend on the type of pore-pore interaction, attractive or repulsive, but are mainly determined by the octagonal symmetry of each single pore. These results pave the way to further studies needed to determine the biological nature of pore-pore interactions.

## 1 Introduction

Genetic information in eukaryotic cells is well protected inside the cell nucleus that is divided from the outside cytoplasm by a membrane called nuclear envelope (NE). This segregation has the advantage of protecting the genome from sources of damage, but on the other hand communications based on exchange of macro-molecules, such as messenger RNAs (mRNA) or transcriptomic factors, are of vital importance during all the cell life cycle, to control protein synthesis and instruct gene expression^1,2^. The spatial architecture of the nucleus is crucial for the interaction between the genome and protein components of the nuclear complex and has a role in chromatin reorganization during cellular differentiation.

Nuclear pores, large protein assemblies inserted in the nuclear envelope, are responsible for selective nucleocytoplasmic transport, allowing the free diffusion of ions and small molecules and acting as selective gates for import and export of macromolecules, such as proteins and mRNAs^3^. Furthermore, nuclear pores are also involved in the organization of the genome and they contribute to gene regulation through physical interactions with chromatin^4^.

Since the discovery of nuclear pores in the 1950s, their structure has been the subject of extensive experimental investigations using Electron Microscopy (EM) and Cryo-Electron Tomography (cryo-ET) ^5^. Great effort was spent on determining the composition of a single pore in terms of different protein complexes. Nuclear pores appear as modular assemblies of discrete constituents arranged with octagonal symmetry around a central axis^6^. Further investigations identified those discrete elements as multiple copies of about 34 protein subunits (nucleoporins). These peculiar proteins are remarkably conserved throughout eukaryotes, showing similar features in algae, yeast, vertebrates such as Xenopus Laevis, up to human.

The characteristic shape of a nuclear pore consists of two superimposed rings of nucleoporins, with eightfold symmetry, one on the outer face of the nuclear membrane and one on the inner face, with eight extended filaments departing from each ring. In contrast, the center of the pore, which forms the permeability barrier, is filled with disordered filaments of phenylanine-glycine (FG) repeats^1,5^. In this context, the experimental observations have already been accompanied by in silico studies involving all atoms and coarse-grained molecular dynamics simulations in order to to characterize the peculiar structure of nuclear pore complexes (NPCs).^7,8,9^.

Despite great progress in understanding the structure of a single nuclear pore, little is known on how nuclear pores are distributed across the surface of the nuclear membrane (in an average human cell there are approximately 2000–3000 nuclear pores) and whether and how they might interact among each other. Interactions are likely modulated by the nuclear lamina, a filamentous protein network underlying the nuclear envelope, but how the interaction occurs is still unclear. Recent observations in Drosophila show that nuclear pores are arranged in a nonrandom manner with clusters that suggest the presence of an effective mutual attractive interaction^10^

Earlier studies, interestingly, revealed that highly proliferative cells such as embryos or tumors have an high density of nuclear pores on the nuclear membrane, while terminal differentiated cells have fewer, suggesting a link between number and distribution of pores and cell activity^11^. This link has been further explored in another early study focused on the changes in distribution of nuclear pores during spermatogenesis, following the evolution from spermatocytes to early spermatids. In particular, a clear change in nuclear pore spatial organization, from aggregation with hexagonal packing in pore rich areas coexisting with large pore-free areas in spermatocytes to a random distribution of pores in early spermatids, has been observed^12^. A further step in our understanding of the role of nuclear pore organization came from the observation of large pore-free islands in HeLa S3 human cells. These islands disperse with cell-cycle progression and reveal the importance of lamin A/C in regulating the pore distribution^13^.

In a recent paper, Selleés et al. performed super-resolution microscopy on Xenopus laevis oocytes observing the variation of nuclear pore distribution on the nuclear membrane during oocyte development^14^. In the present paper, we analyze those experimental data^14^ to investigate the spatial distributions of nuclear pores across the nuclear membrane during the development of Xenopus laevis oocytes. To this end, we use tools typical of soft matter physics, such as the radial distribution function (RDF), local order parameters and Voronoi tessellation. To model the spatial distribution of nuclear pores observed experimentally, we introduce an effective interaction among them. We first define a potential with octagonal symmetry to properly model the shape of nuclear pores and then perform extensive numerical simulations of interacting nuclear pores, studying their behavior as the density varies. So far attempts of modeling NPCs geometrical arrangement on the nuclear surface and using computer simulations to deepen our understanding on it were missing. With our work we try to fill that gap proposing a first attempt to study in silico peculiarities and features of the spatial distribution of nuclear pores.

## 2 Materials and Methods

### 2.1 Experimental images

In this Section we analyze super-resolution experimental images of nuclear pores of *Xenopus laevis* oocyte by Sellés et al. ^14^. According to the stage of development of the oocyte^15^ we identify three groups of images: an early Stage II, an intermediate Stage IV, and a later Stage VI. A different number of samples were taken at each stage, specifically: 6 samples for Stage II, and 11 samples for both Stage IV and Stage VI. The images are 2560 × 2560 pixels (px) wide, with 1 px corresponding to 10 nm. Examples of nuclear pore experimental images are reported in Fig. 1a)-c).

**Fig. 1.**
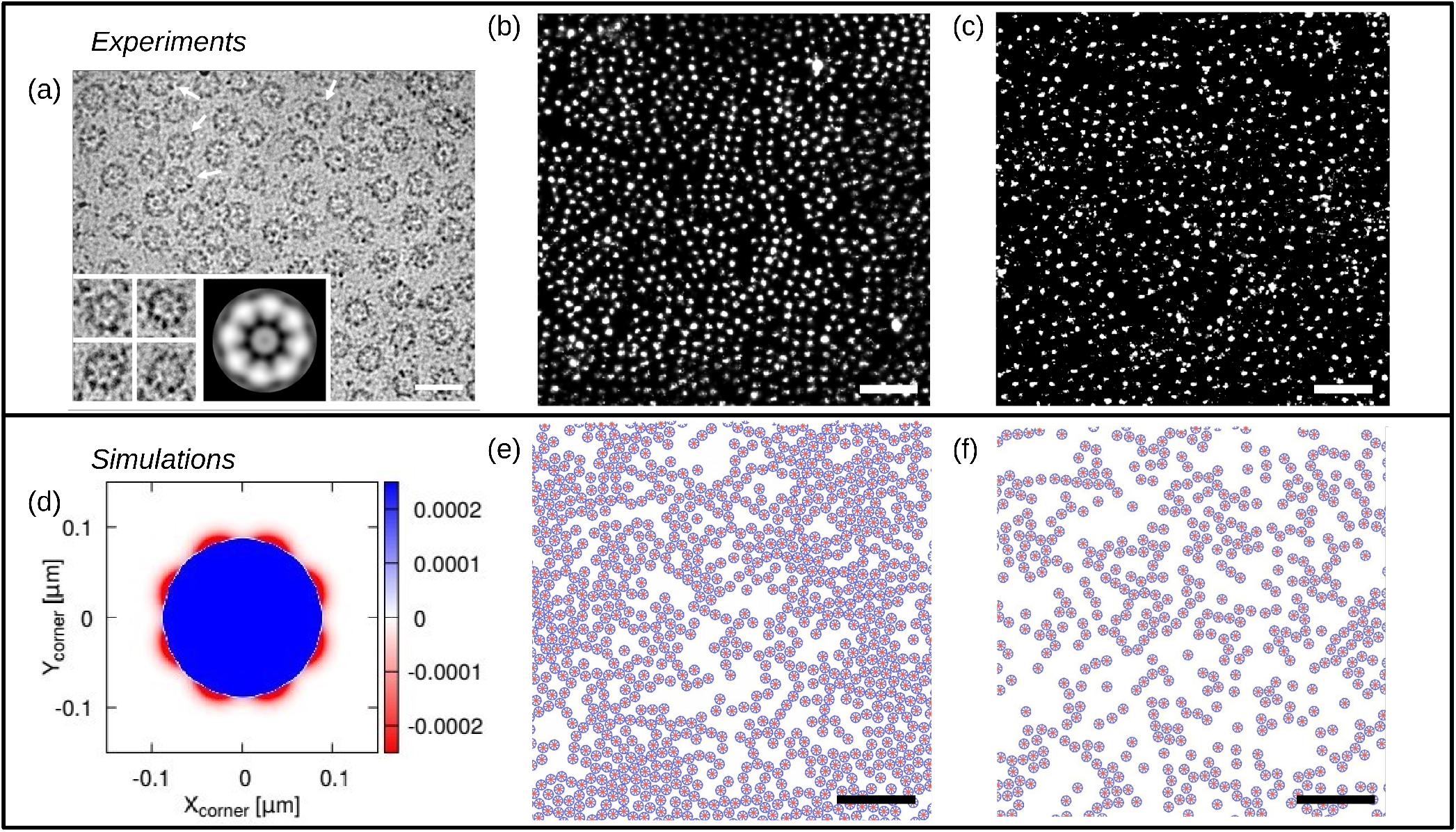
Comparison of experimental and simulated nuclear pore images. (a) Energy-filtering transmission electron microscopy (EFTEM) images of Xenopus oocyte NEs embedded in thick amorphous ice. From those images the eight-fold symmetry of nuclear pores can be appreciated. The four insets on the bottom left of the figure show higher magnification images. Correlation averages over 100 pores are plotted in the bottom right inset, clearly revealing the shape of a single pore. Scale bar represents 200 nm. Reprinted from ‘Cryo-electron tomography provides novel insights into nuclear pore architecture: implications for nucleocytoplasmic transport’ Daniel Stoffler et al., Journal of molecular biology, 2003, 328.1: 119-130.^21^, Copyright 2003, with permission from Elsevier. (b),(c) Portions of experimental images of nuclear pores in *Xenopus laevis* oocyte at different developmental stages, respectively Stage II (b) and Stage VI (c), obtained using super-resolution microscopy. Scalebars are 1 *μ*m. Panels (b) and (c) are adaptations from Sellés et al. ^14^. Scalebars are 1 *μ*m. (d) Potential energy surface obtained from the modeled interaction between a pore and a corner of a neighboring pore: the blue area marks a strongly repulsive region, while the red areas mark the trapping centers. (e),(f) Example of two configurations of nuclear pores obtained from numerical simulations for comparison with experimental data. The density is 36 NPC/*μ*m^2^ for (e) and 20 NPC/*μ*m^2^ for (f). Scalebars are 1 *μ*m.

#### 2.1.1 Tracking of Nuclear Pores

We analyze the trajectories of the nuclear pores with Trackpy v0.4.2, a Python package for particle tracking in 2D, 3D, and higher dimensions^16^. In particular, we first discriminate the nuclear pores using the function *trackpy.locate,* whose working principle is the following: i) preprocess the image by applying a bandpass filter (i.e. performing a convolution with a Gaussian to remove short-wavelength noise, and subtracting out long-wavelength variations by subtracting a running average, in order to retain intermediate scale features), ii) apply a threshold over the color channels, and iii) locate all the peaks of brightness, each referring to the position of a pore^16,17^. The parameters used for the tracking are: *diameter* = 9 px the diameter, and *minmass*, the minimum integrated brightness, working as a threshold value. The latter value is chosen based on the samples: *minmass* = 0 (no threshold) is used in the high-density samples of Stage II, while higher values of this parameter were necessary to correctly detect pores in the noisier experimental images of Stage IV and VI. The tracking procedure also allows us to determine the density of pores defined as the number of pores per unit area of the nuclear envelope from the experimental samples: 34.9 ± 2.3 NPC/*μ*m^2^ for Stage II, 25.6± 2.3 NPC/*μ*m^2^ for Stage IV, 20.5 ± 1.7 NPC/*μ*m^2^ for Stage VI. The errors here represent the standard deviation computed on the ensemble of samples for each developmental Stage. The above density values are slightly lower than those computed by Sellés et al. ^14^. We attribute this discrepancy to the different tracking techniques employed and to an inherent uncertainty related to experimental measurements performed with super-resolution optical microscopy. In fact, with this technique, some fluorescent spots may be “fragmented” in the final image, due to small microscope movements. Thus it can happen that a single pore appears divided into several dots, introducing a certain arbitrariness in the counting of pores.

### 2.2 Numerical Simulations

To model the interaction among nuclear pores, we consider their peculiar octagonal shape, as observed in early experimental studies^6,18^ (see e.g. Fig. 1a) and subsequently confirmed by structural studies on nucleoporins, and by recent advances in experimental techniques such as cross-linking mass spectrometry and cryo–electron tomography^5^. Coarsegrained models are of key importance for understanding the essential behaviors of biological phenomena without resorting to detailed modeling of the molecular structure^19^. In the following paragraph, we describe the coarse-grained model providing the details on the form of the potential for the pore–pore interaction, and the simulation protocol.

#### 2.2.1 Model Potential for Nuclear Pores

To account for the composite structure of each nuclear pore and its overall octagonal shape, we implement a coarse-grained model of the pore, which consists of a central particle surrounded by eight particles located at the vertices of a regular octagon, of circumradius *R*. For simplicity, the pore is treated as a rigid non-deformable object. Taking experimental data as reference^14,20^, in all simulations we set *R* = 67.5 nm. The overall interaction potential acting among the particles of two neighboring pores is composed of three terms, each of which consists of a Lennard-Jones (LJ) potential,

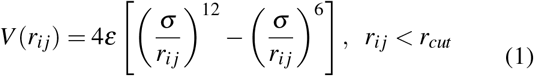

with *r_ij_* = |**r***_i_* – **r***_j_*| is the distance between particle *i* and particle *j,* and *ε, σ*, and *r_cut_* parameters depend on the interaction term:

- **center-center interaction** – the central particle of a pore interacts repulsively with the central particle of a neighboring pore. For this term we set *ε_cc_* = 0.01 pg·*μ*m^2^/*μ*s^2^, *σ_cc_* = 0.12 *μ*m, and the cutoff distance is set 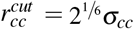, to make the interaction purely repulsive. This term is necessary to avoid non-physical configurations, such as the case of overlapping pores that are otherwise rarely encountered.
- **center-vertex interaction** – the central particle of a pore interacts repulsively with the particle at the vertex of a neighboring pore. Again, this is introduced to avoid pore overlap and interpenetration. LJ parameters for this term are *ε_cv_* = 0.01 pg·*μ*m^2^/*μ*s^2^, *σ_cv_* = 0.08 *μ*m, 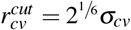.
- **vertex-vertex interaction** – the particle at the vertex of a pore interacts with the particle at the vertex of a neighboring pore. We first considered the full LJ interaction (long-range attractive, short-range repulsive), but the purely repulsive case was also studied (see Supplementary Fig. S1). For the LJ interaction we set *ε_vv_* = 5·10^-4^ pg·*μ*m^2^/*μ*s^2^, *σ_vv_* = 0.02 *μ*m, and the cutoff distance is set to 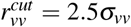 so to include the attractive part. For the purely repulsive case we set 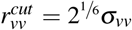. By construction, in our model the overall pore–pore interaction is mainly driven by the present term, due to the fact that the central particles are surrounded by vertex particles.

In all cases, the LJ potentials are shifted to zero at the cutoff distance to avoid any energy discontinuity. Fig. 1d reports the total interaction energy of an octagonal pore (centered at the origin) with a corner particle from a second pore, as a function of the latter’s position. The resultant potential energy surface (PES) shows a central strongly repulsive region (in blue) and trapping regions (in red) concentrated near the eight corner sites.

#### 2.2.2

In silico nuclear pores configurations

The simulations of nuclear pore assemblies are performed using LAMMPS^22^ with a timestep Δ*t* = 10^-5^ *μ*s. The starting configurations consist of 1000 randomly placed octagonal pores confined in a square periodic box of side *L* = 40 *μ*m. To mimic the different experimental nuclear pore densities observed during oocyte development, the random configurations are alternately subjected to 10^5^ steps of box compression at a constant temperature 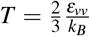 followed by a 2·10^5^ steps of annealing from high temperature 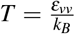 down to 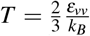, in order to allow thermally-assisted rearrangements. Such procedure is iterated until the required density is obtained, and a final energy minimization at *T* = 0 K in 2× 10^6^ steps is performed. Following the above protocol we obtain configurations with a density of 20, 26, 36, 46, and 53 NPC/*μ*m^2^. For each density value we obtain 10 different realizations starting from different random initial positions, in order to allow for proper statistical averaging. Examples of nuclear pore configurations obtained with numerical simulations are shown in Fig. 1e) and f).

### 2.3 Statistical analysis of nuclear pores structure

In this Section we describe three different quantities used to provide a statistical comparison of the simulations with the reference experimental data: the radial distribution function, the hexatic order parameter and voronoi diagram.

#### 2.3.1 Radial Distribution Function

To gain insight on the local structure of the nuclear pore complex on the nuclear membrane, we analyze the radial distribution function (RDF)^23^:

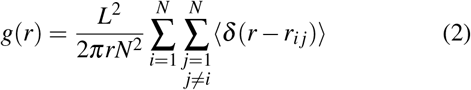

where *N* is the number of particles in the system, *L* is the system size as specified above, and *r_ij_* is the distance between particles *i* and *j* and the average 〈〉 is over particles.

The RDF is a key tool for the theory of monoatomic liquids, to characterize amorphous colloidal solids^24^ and to study glasses and the glass transition^25^.

#### 2.3.2 Hexatic Order Parameter

To characterize the geometrical properties of the structure formed by the nuclear pores, we compute for each particle the *n*-fold local orientational order parameter (hexatic order parameter):

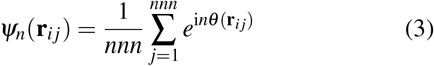

where *nnn* is the number of nearest neighbors of particle *i*, *θ*(**r***_i j_*) is the angle formed by the *x* axis and the vector **r***_i j_* connecting particles *i* and *j*. Experimental nuclear pore complexes of *Xenopus laevis* at different developmental stages show ordered regions characterized by triangular and square lattice^14^, therefore we focus our analysis on order parameters i) with *n* = 6 for which |*ψ*_6_| = 1 for particles belonging to a perfect hexagonal structure, and ii) with *n* = 4 for which |*ψ*_4_| = 1 for particles belonging to a perfect square lattice. The determination of nearest neighbor particles is performed using a cutoff distance *σ_cut_* = 0.15 *μ*m for the simulations and *σ_cut_* = 0.20 *μ*m for the experimental images. Those values have been chosen considering the typical inter-particle distance in simulated and experimental samples. For all isolated particles (nnn < 2) we set *ψ_n_*=0.

#### 2.3.3 Voronoi Tessellation

We finally analyze the Voronoi diagram for the nuclear pore configurations, partitioning the image into regions of convex polygons around the center of each pore. This so-called Voronoi cells represent the area of space containing all points that are closer to one pore than to any other. For the experimental images, the coordinates of the pore centers were obtained from the tracking analysis and used as input for the Voronoi tesselation. For the simulations only the central particle of each octagon is considered in the Voronoi analysis. By construction, each Voronoi cell has polygonal shape, with a number of sides that corresponds to the number of neighbors. To compute the Voronoi tesselation we used the Python library Freud^26^, that allows to account for periodic boundary conditions. Using this method we extract for each pore the number of neighbors (pores are considered neighbors if they share an edge in the Voronoi diagram) and the size of each associated Voronoi cell.

## 3 Results

### 3.1 Global structure of NPC

From the RDF of the experimental samples, we note that at high density (early stage of development of the oocyte) *g*(*r*) shows a liquid-like shape, with two peaks clearly visible (see Fig. 2a). In fact in monoatomic liquids, at short range *g*(*r*) shows a pattern of peaks representing the nearest neighbour distances, and at large *r* it tends to unity due to total loss of order^23^. As the density decreases during oocyte development (Fig. 2b,c), the second peak of the experimental *g*(*r*) tends to disappear while the first peak tends to flatten out, thus converging toward a gas-like phase in which the order is lost. Previous analysis of the experimental images suggested a significant presence of square lattice domains of nuclear pores at low density (Stage II)^14^. For this reason, we have looked for a specific peak in the *g*(*r*) function. In the case of a regular square lattice, a second peak should be present at 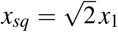 (where *x*_1_ represent the spatial coordinate of the first peak of the *g*(*r*) function), however we notice that a peak in this position is not visible. Nevertheless *x_sq_* falls on the tail of the first peak (which is quite flat), suggesting that some sparse regions with square lattice could be present inside the amorphous liquid. Similar considerations have been done for hexagonal structures at high densities: if a regular hexagonal packing of pores is present, the *g*(*r*) should display a peak at the position 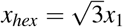 and again, this is not the case, with *x_hex_* falling only on the growing part of the second peak for the high density configurations. Some hexagonal structures at high density could be present but are not predominant, being the spatial distribution of NPCs mainly disordered. The *g*(*r*) obtained from simulations with the octagonal attractive potential (Fig. 2) reproduces the above described liquid-like profile presenting just a few peaks emerging over an otherwise flat profile. Furthermore, the positions of the peaks in the simulated *g*(*r*) match nicely with the experiments (especially for what concerns the first peak), suggesting that our model is able to capture the relevant features of the effective interaction among the nuclear pores on the nuclear membrane. This is a non-trivial result, as the position of the *g*(*r*) peaks closely reflects the interactions held between the constituents, and are a signature of each material and its peculiar properties as shown in previous studies of noble gases or water^27–30^. Ultimately, from this analysis, we can rule out the significant presence of extensive crystalline regions.

**Fig. 2.**
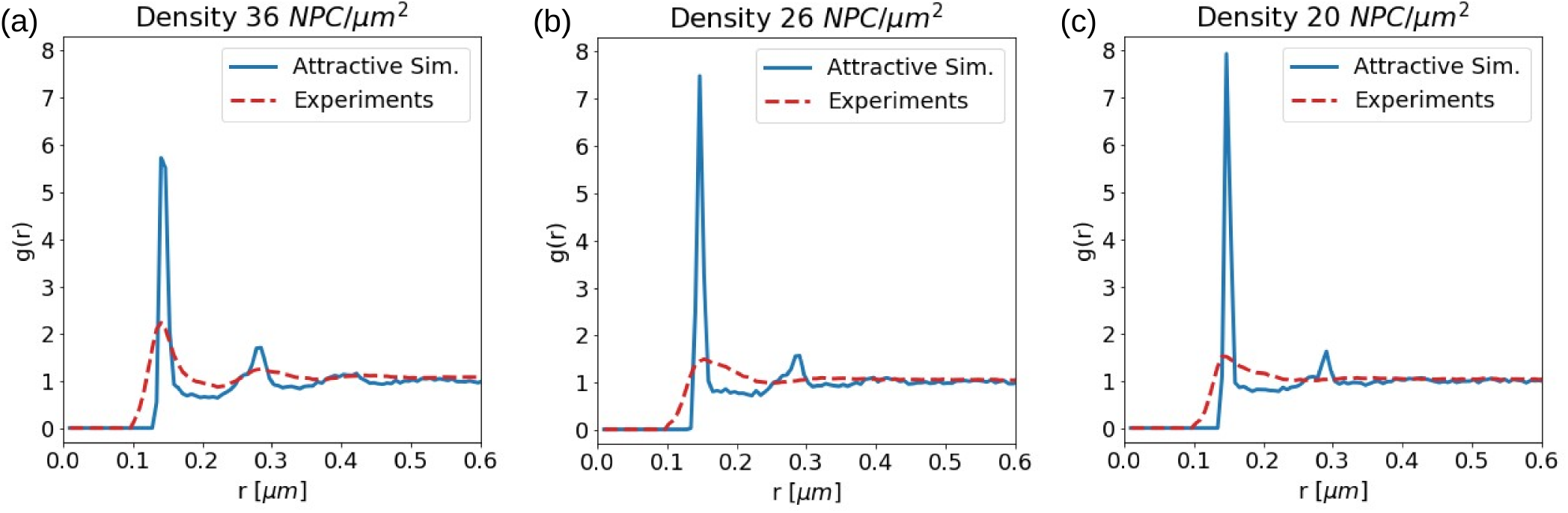
Radial Distribution Function of nuclear pores. The *g*(*r*) is reported for three different densities of the nuclear pores. Blue curves are from averages over ten simulations with attractive octagon potential. Dashed red curves are obtained from the experimental images for Stage II - (a), Stage IV - (b), Stage VI - (c).

### 3.2 Orientational Order

In Fig. 3, we report the calculated local order parameter *ψ*_6_ and *ψ*_4_, as defined in Section 4, for some experimental samples and for the configurations obtained from simulations. The cutoff to consider a pore as a neighbor was set to *σ_cut_* = 0.20 *μ*m for the experiments and *σ_cut_* = 0.15 *μ*m for the simulations. These values were chosen considering the typical pore-pore distances. From Fig. 3a and Fig. 3c it can be observed that only few pores are associated with square symmetry (*ψ*_4_ ~ 1). Instead, Fig. 3b and Fig. 3d show much broader regions associated with triangular lattice structures (*ψ*_6_ ~ 1). In those samples, in presence of higher pore densities, more nuclear pores belong to a regular structure and some regions with clear hexagonal order appears. To further investigate such behavior we computed the distribution of the local order parameters *P*(*ψ*_6_) and *P*(*ψ*_4_) averaged over all the samples for each value of the density (see Fig. 4). First, we can observe that the distributions of the experimental samples (Fig. 4a,b) are all unimodal, thus confuting the hypothesis of two different coexisting phases suggested previously^14^. Secondly, we notice that for both *ψ*_6_ and *ψ*_4_ the distributions are peaked at a value which increases with the density. The largest mode value is obtained for *ψ*_6_ at the highest density (Stage II), suggesting a preference for hexagonal structures in the dense limit. The above trends of the distributions with the density are well reproduced by the simulated NPCs, as reported in Fig. 4c and Fig. 4d. For a more straightforward comparison we have reported in Fig. 4e the average value of |*ψ*_6_| and |*ψ*_4_| as a function of the pores density for both simulations and experiments. For the former, we observe that both |*ψ*_6_| and |*ψ*_4_| values slowly increase with the density, showing the same values until a density of 36 NPC/*μ*m^2^ where a bifurcation occurs. Beyond that density value, |*ψ*_4_| seems to saturate while |*ψ*_6_| keeps increasing, thus favoring the hexagonal order at high density. It is worth to note that, in the explored density range, the |*ψ*_6_| value is far from approaching unity, corresponding to a crystalline structure, indicating that much larger densities would be required for such an ordered phase. Finally we note that in Fig. 4e the points associated to the experimental data do not fall exactly on the theoretical curves derived from the simulations. This can be partially explained with the uncertainties connected with the experimental observations of the nuclear pores, that affect the density evaluation. Despite that, the experimental points show a trend very close to that of the simulation, with an initial overlapping of |*ψ*_6_| and |*ψ*_4_| values, and a further bifurcation at higher density.

**Fig. 3.**
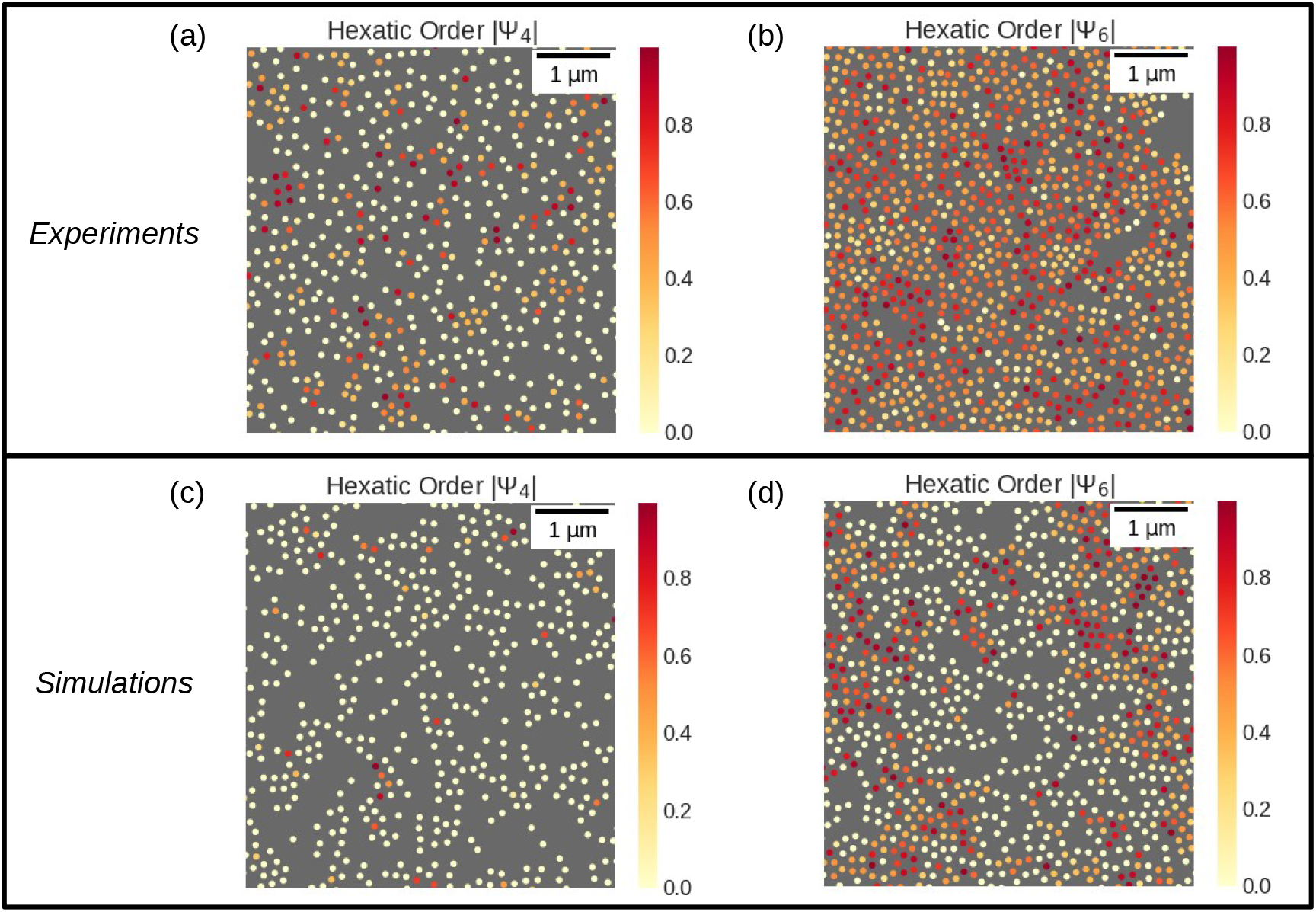
Color maps of local order parameters. Snapshots of pores colored as a function of the local order parameter: (a) a zoomed region of an experimental sample at Stage VI, for which we estimate a density of 20.5 ± 1.7 NPC/*μ*m^2^; (b) a zoomed region of an experimental sample at Stage II, for which we estimate a density of 34.9 ± 2.3 NPC/*μ*m^2^; (c) and (d) a simulation box with pores density 20 NPC/*μ*m^2^ and 36 NPC/*μ*m^2^ respectively.

**Fig. 4.**
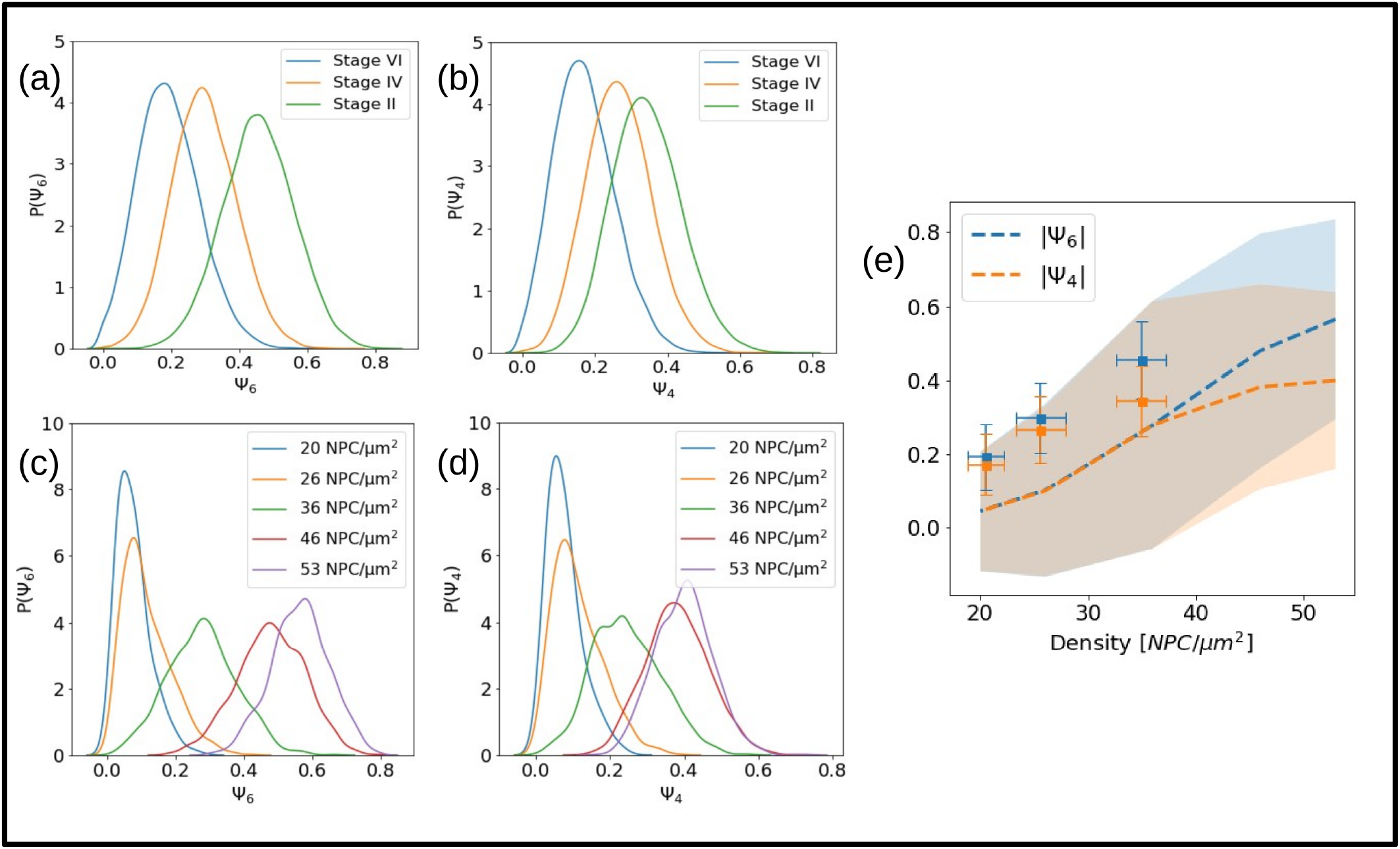
Distribution of the local order parameters. The distribution of the local order parameters at different pores densities is reported for (a),(b) experimental samples and (c),(d) for simulated configurations; (e) the average |*ψ*_6_| and |*ψ*_4_| values as a function of the density. The dashed lines report the values obtained from the simulations, with shadows highlighting the respective standard deviation. The points represent the values computed from the experimental samples, with vertical errorbars for the standard deviation, and horizontal errorbars reporting the error on the density, as described in Section 2.1.1.

### 3.3 Properties of Voronoi cells

Examples of Voronoi tesselation performed on high density NPC from the experimental images and from the simulated configurations are reported in Fig. 5a and Fig. 5d, respectively. The comparison in these high density samples shows similar tessellation patterns in experiments and simulations. A statistical analysis on the number of sides *N* of the Voronoi cells at different densities (Fig. 5b and Fig. 5e) clearly shows that the *N* = 6 occurrence increases with the density, and viceversa for the the *N* = 4 occurrence. Therefore, the Voronoi analysis enforces the idea, already anticipated above by the local order parameter, that the hexagonal configuration is favored at high densities at the expense of other kind of local order arrangements. Correspondingly, in agreement with experimental observations, the presence of some square structures at low density is also supported. However, we note at any density a significant number of cells with *N* = 5, about half between those with *N* = 4 and *N* = 6. We associate this with the particular sensitivity of the Voronoi tesselation method to “defect”, i.e. deviations with respect to the ideal symmetric cases. To provide a further comparison, we report in Fig. 5c and Fig. 5f the distribution of the Voronoi cells area. Again, a good agreement between the experiments and our model is obtained, with much narrower distributions at higher densities, shifting toward higher area values and broadening out as density decreases.

**Fig. 5.**
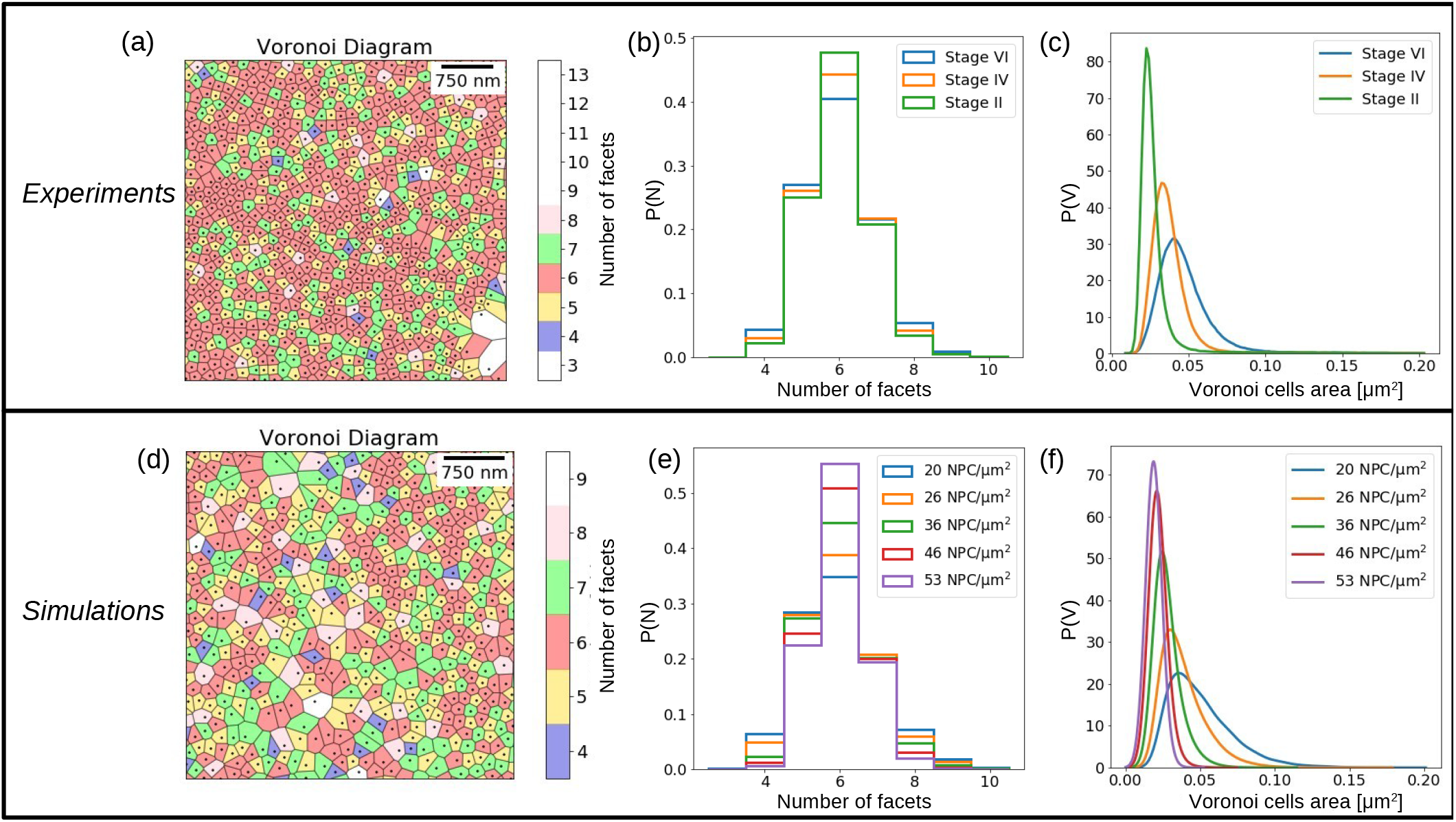
Voronoi tesselation applied to nuclear pores. Examples of Voronoi tesselation are provided for (a) an experimental samples at Stage II and (d) for a simulation with density 36 NPC/*μ*m^2^; (b) and (e) the histogram of the number of Voronoi cell facets, for different densities; (c) and (d) the corresponding distribution of the Voronoi cells area.

## 4 Discussion and Conclusions

A first attempt to investigate the spatial distribution of nuclear pores goes back to the ‘70s, when the positions of NPCs on the surface of rat kidney nuclei was observed and distances among them measured ^31^. Already in this study, some regularities were found in the distribution of pore-pore distances measured in the samples, suggesting a non random spatial distribution and some peaks corresponding to hexagonal structures, even if the statistics was too poor to reach further conclusions. More recently, Selleés et al. investigated the angular distribution between first neighbors of nuclear pores, revealing no preferential angles for Stage II and IV *Xenopus laevis* oocytes (high density) and two distinct peaks at 90° and 180° for later Stage VI, suggesting the presence of square lattice regions at low density^14^. In our study, from the analysis of the radial distribution function *g*(*r*) of the nuclear pores on the nuclear membrane of *Xenopus laevis* oocytes, we could not observe peaks in correspondence of peculiar geometrical structures, meaning that even though some crystalline regions are present, they are quite rare and do not statistically influence the overall NPC spatial distribution. Interestingly by analysing the *g*(*r*) of the nuclear pores, we were able to identify an amorphous, liquid-like structure in which, in the early phase of oocyte development (when NPC density is high), long-range order is soon lost. On the other hand, as the oocyte develops, the nuclear pore density decreases and *g*(*r*) shows a behaviour compatible with a more dilute, gas-like system. From a biological point of view, the early stages of oocyte development are associated with intense transcriptional activity, as the oocyte needs to build up a huge reserve of gene products such as mRNAs, tRNAs and proteins in order to correctly fulfil its future role after fertilisation. Once the necessary maternal mRNAs have been copied, transcriptional activity in the later stages of oocyte development becomes lower ^32^. These changes in transcriptional activity could be linked to changes in the spatial distribution of NPCs during oocyte development, particularly changes in density. It would be extremely interesting to further explore this connection from a biophysical point of view, e.g. by trying to quantify the flow of matter through the pores, (as has already been done in some previous kinetic studies, which showed that a single NPC can allow a mass flow of nearly 100 MDa/s^33^), at different stages of the cell’s life-cycle. The positions of the peaks in the *g*(*r*) we computed for nuclear pores are another key point of our results. Indeed, it is known from the physics of matter that the positions of the peaks of the radial distribution function and their relative distances give actual information about the geometrical arrangement of the particles within a material, and are a signature of the material itself ^34^. In particular, the peak positions allow to indirectly infer the type of interactions among the constituents of a specific material. Here, we have shown that the eight-fold potential used to model the NPC in our simulations is able to nicely reproduce the experimental *g*(*r*) peak locations. In particular, the position of the first peak obtained from the simulations is in excellent agreement with Sellés et al. ^14^ and with previous observations^12,31^. We also checked that assuming a simple LJ potential acting among the pores (i.e. with spherical symmetry) with parameters compatible with experimental pore sizes, the prediction of the *g*(*r*) peaks is not in agreement with experimental results (see Supplementary information). Even if the hypothesis of a pore-pore potential with spherical symmetry (perhaps with an effective interaction size for the pore that does not coincide with its physical size) can not be excluded, our work suggests that the octagonal shape of the pore and the associated eight-fold symmetry of its interaction potential plays a crucial role in determining the correct spatial distribution of the pores. These facts are worth to notice, since our simplified model based on the assumption of an effective eight-fold pore-pore interaction is able to catch a crucial signature of the spatial distributions of nuclear pores, the radial distribution function peaks positions, even if the interaction details (e.g. if the pore-pore potential is attractive or repulsive) are not known (see Supplementary). Hopefully this could help to deepen the investigation of the nature of the pore-pore interplay, allowing to study also *in silico* an interaction that in reality is not fully understood under a biological point of view. Pore-pore interactions are unlikely to be direct, but rather mediated by the lamin scaffold through complex interactions that are hard to model explicitly. In this sense, our assumption that nuclear pore arrangement can be modeled as assembly of interacting octagons is possibly oversimplified, since after development pores are stuck within the lamina and are unlikely to diffuse. We can, however, imagine that as the nuclear envelope is formed the nuclear pores are arranged in a way that is dictated by their geometry and which could then be captured by our model. Since extensive MD simulations of lamina filaments forming a three-dimensional network beneath the nuclear envelope have recently been performed^35^, it would be interesting to try to go further in modelling the outer regions of the cell nucleus, linking the lamina network and the spatial distribution of the nuclear pores.

Coming back to our analysis, the spatial distribution of NPCs, investigated through the local order parameters shows that at high density the pores tend to arrange following the triangular lattice. Even though the *g*(*r*) does not show explicit peaks in correspondence to a triangular lattice, the study of *ψ*_6_ (Fig. 3) and the Voronoi tessellation method (Fig. 5) prove that at high density islands of six-fold symmetry packed pores appear. Noticeably, such behavior has been already reported in previous experimental observations. During apoptosis the distribution of nuclear pores on the cell nucleus strongly changes, bringing the NPCs to be highly concentrated in small regions of the nuclear envelope (on mouse cell nuclei) and leaving the rest of the surface pore-free. Those clusters of pores showed a hexagonal packing and were supposed to be correlated with diffuse chromatine areas^36^. Occasional areas of very regular hexagonal packing of nuclear pores have been also observed to emerge during the development of male germ cells, in rodent spermatocytes^12^. Those facts open interesting questions on how the geometrical disposition of the pores in some areas, or even more simply, their density, are influenced by the underlying nuclear activity and on what are the biological causes responsible for the effective interaction among NPCs. Considering the pores under a geometrical and topological point of view, underlying the importance of their octagonal shape, like our simple model does, could be extremely interesting also in the contest of membranes studies. In fact in a recent paper by Torbati et al.^37^, the authors studied the mechanical stability of the lipid bilayer membrane of the nuclear envelope, considered as two concentric membrane shells fused at numerous sites with toroid-shaped nuclear pores (here simply modeled as circular holes). Using mechanistic arguments based on elasticity, they showed that in- and out-of-plane stresses can give rise to the pore geometry and the geometric topology observed in cell nuclei, finding simulated interpore distances in good accord with the ones observed in mammalian cells nuclei. How octagons can contribute to stabilize the curvature of a spherical membrane^1^ and how they tend to be spatially arranged on such a geometry could be an issue to consider to better clarify the process of nuclear pores formation.

## Acknowledgements

We thank Zoe Budrikis for preliminary developments of the simulation code.

