## Supplementary figures for "Spatial organization of nuclear pores in *Xenopus laevis* oocytes" (from another version of this preprint: version 3, 2021-11-24)

### Spatial organization of nuclear pores in Xenopus laevis oocytes : supplemental document

#### 1 Introduction

In this supplementary file some results obtained using alternative potentials for pore-pore interactions are showed. Those results have been compared with the eight-fold potential with attractive corners that represents the standard model used to obtain all the results showed in the paper.

#### 2 Alternative models for pore-pore interaction

Two alternative potential for pore-pore interaction have been tested.

- eight-fold potential fully repulsive
- simple Lennard-Jones (spherical simmetry)

The repulsive octagonal potential is obtained using the same LJ parameters of the attractive potential used in the paper, except for the vertex-vertex interaction in which is truncated to be fully repulsive: $r^{cut}_{vert-vert} = 2^{\frac{1}{6}}\sigma_{vert-vert}$. The simple spherical potential is obtained with the following LJ parameters: $\epsilon = 5 \cdot 10^{-4}\,\mathrm{pg}{\cdot}\mu\mathrm{m}^2/\mu\mathrm{s}^2$, $\sigma = 0.12\,\mu\mathrm{m}$, $r_{cut} = 2^{\frac{1}{6}}\sigma$. In Figure S1 the potential energy surfaces of the two alternative potential studied are shown.

The statistical analysis methods described in the paper (see Material and Methods) have been applied to characterize the distributions of nuclear pores obtained assuming those alternative pore-pore potentials. In Figure S2 the radial distribution function of the pores distributions obtained with different models are illustrated. What emerges is that both the repulsive and attractive eight-fold potentials seem to describe quite well the experimental peak location, in particular for what concerns the first peak of the $g(r)$ (see Figure S2 c). The potential with spherical symmetry, instead, does not match the experimental peaks. As already mentioned in the paper, we have assumed a LJ potential acting among the pores with parameters compatible with experimental pore sizes ($\sigma = 0.12\mu m$). For this reason, the hypothesis of a pore-pore potential with spherical symmetry can not be completely excluded, perhaps considering an effective interaction size for the pore that does not coincide with its physical

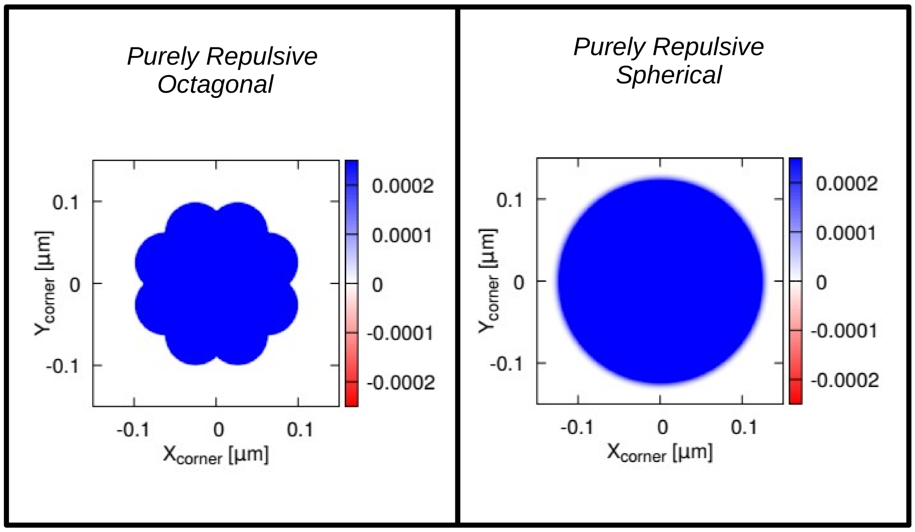


Fig. S1: Potential energy surface obtained from the modeled interaction between a pore and a corner of a neighboring pore: blu area marks strongly repulsive regions. (a) represents the eight-fold fully repulsive potential, (b) the LJ potential.

size. Despite from that, our work seems to suggest that the octagonal shape of the pore and the associated eight-fold symmetry of its interaction potential plays a crucial role in determining the correct spatial distribution of the pores, no matter about the attractive or repulsive nature of the pore-pore interaction.

Furthermore, the local order parameters described in the paper (see Material and Methods) have been computed also for the pores configurations obtained with the alternative potentials. In Figure S3 the value of $|\Psi_4|$ and $|\Psi_6|$ averaged over particles and configurations have been plotted for different pore densities. A substantially different behavior is observed for the pore-pore interaction with spherical symmetry, compared to the values of the local order parameter obtained from the reference eight-fold attractive potential. The simple LJ potential gives a trend that is different from the one that the experimental data suggest: $|\Psi_6|$ is always bigger that $|\Psi_4|$, even at small densities, correctly revealing the preference of spheres for hexagonal packing. Again, also for what concerns the local order parameters, no significant differences have been reported for the eight-fold repulsive potential respect to the eight-fold attractive potential.

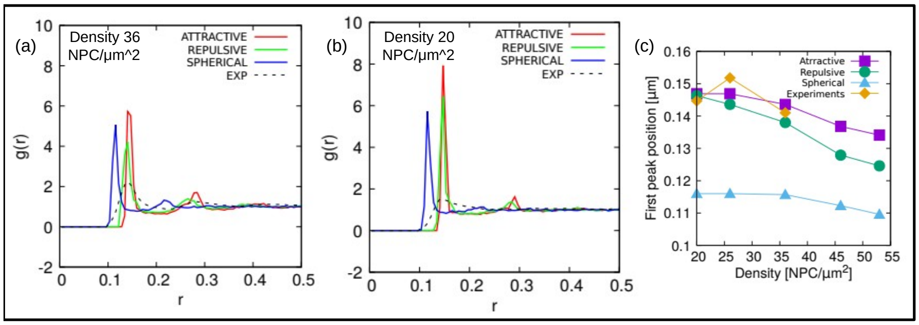


Fig. S2: **Alternative models for pore-pore interaction.** – (a) The radial distribution function for different interaction potential: attractive octagonal potential (red curve), repulsive octagonal potential (green curve) and simply spherical repulsive potential (blue curve). Data are relative to simulations with a density of $36\,\mathrm{NPC}/\mu\mathrm{m}^2$, and compared with the $g(r)$ from experimental images of Stage II oocytes (black dashed line); (b) the same results are shown for simulated densities of $20\,\mathrm{NPC}/\mu\mathrm{m}^2$ and experimental samples on stage VI; (c) the position of the first $g(r)$ peak as a function of the density.

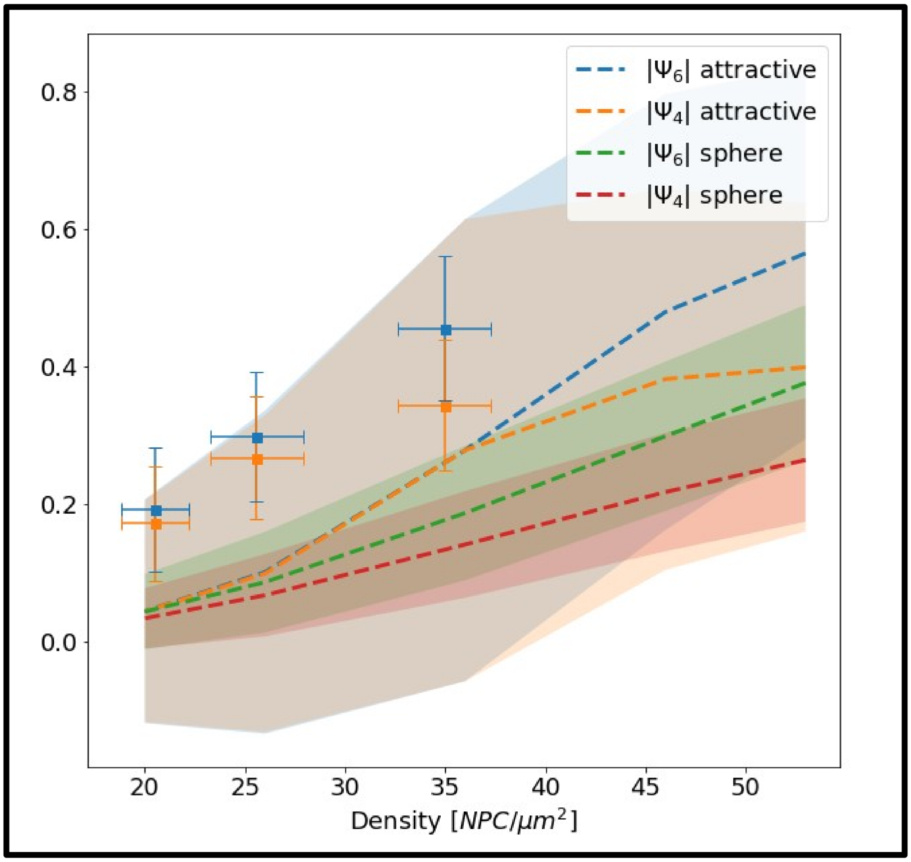


Fig. S3: **Local order parameters for alternative pore-pore interactions** – The average $|\psi_6|$ and $|\psi_4|$ values as a function of the density for different pore-pore interactions. The dashed lines report the values obtained from the simulations, with shadows highlighting the respective standard deviation. In blue and orange can be observed the local order parameters for the reference eight-fold attractive potential, while the dashed red and green curves are referred to the LJ pore-pore interaction, with simple spherical symmetry. The points represent the values computed from the experimental samples, with vertical errorbars for the standard deviation, and horizontal errorbars reporting the error on the density.
